# On Transformative Adaptive Activation Functions in Neural Networks for Gene Expression Inference

**DOI:** 10.1101/587287

**Authors:** Vladimír Kunc, Jiří Kléma

**Affiliations:** Department of Computer Science, Czech Technical University in Prague, Prague 121 35, Czech Republic

## Abstract

**Motivation:** Gene expression profiling was made cheaper by the NIH LINCS program that profiles only ~1, 000 selected landmark genes and uses them to reconstruct the whole profile. The D–GEX method employs neural networks to infer the whole profile. However, the original D–GEX can be further significantly improved.

**Results:** We have analyzed the D–GEX method and determined that the inference can be improved using a logistic sigmoid activation function instead of the hyperbolic tangent. Moreover, we propose a novel transformative adaptive activation function that improves the gene expression inference even further and which generalizes several existing adaptive activation functions. Our improved neural network achieves average mean absolute error of 0.1340 which is a significant improvement over our reimplementation of the original D–GEX which achieves average mean absolute error 0.1637

## 1 Introduction

Gene expression profiling is a great tool for medical diagnosis and deepening of disease understanding (e.g., [1]–[4]). Despite a significant price drop in recent years, the gene expression profiling is still too expensive for running large scale experiments. In order to economically facilitate such experiments, the LINCS program resulted in the development of the L1000 Luminex bead technology that measures the expression profile of ~1,000 selected *landmark genes* and then reconstructs the full gene profile of ~10,000 *target genes* [5]. The original method used the linear regression for the profile reconstruction due to its simplicity and scalability, which was then improved by a deep learning method for gene expression inference called D–GEX [6] which allows for reconstruction of non-linear patterns.

The D–GEX is a family of several similar neural networks with varying complexity in terms of a number of parameters used for the gene expression inference. Our main contribution is that the D–GEX family performance can be further improved by using more suitable activation functions even while keeping the architecture without changes.

We first briefly present the use of artificial neural networks (ANNs) in biology and the usage of adaptive activation functions (AAFs) for improving performance of ANNs. Then we describe used data and their preprocessing before we introduce the original D–GEX and our novel adaptive activation function. We also present several experiments to show that our proposed transformative adaptive activation function (TAAF) can significantly improve the performance of the original D–GEX.

## 2 Related works

### 2.1 Artificial neural networks in biology

Artificial neural networks (ANN) represent a state-of-the-art approach in many fields (e. g. image classification, segmentation or reconstruction, natural language processing or time-series forecasts) and biology is not an exception (review e. g. [7]–[9]). One of such applications is D–GEX [6] which infers a full gene profile using only ~1,000 selected *landmark genes*. The D–GEX family is made up of 9 different architectures. Due to technical reasons, the D–GEX consists of two separate feedforward NNs having from one to three hidden layers each having either 3,000, 6,000, or 9,000 neurons. Each network predicts only a half of target genes (~4,760 genes) and is trained on a separate GPU. The NNs were trained using a standard back-propagation algorithm with mini-batch gradient descent with momentum and learning rate decay [6]. The initial weights were initialized using normalized initialization [10]. The used error metric was *mean absolute error* (MAE).

The original D–GEX was evaluated using data from three different sources — *GEO expression data* curated by the Broad Institute, *GTEx expression data* consisting of 2,921 gene expression profiles obtained using Illumina RNA–Seq platform [11] and *1000 Genomes expression data* consisting of 462 gene expression profiles obtained also using Illumina RNA-Seq platform [12]. The *GEO expression data* contained biological or technical replicates, the final dataset contained ~110,000 samples after their removal. All three datasets were jointly quantile normalized and then standardized for each gene individually.

The D–GEX NNs were compared with linear regression and k–nearest neighbor (KNN) regression. The linear regression builds a linear model for each target gene while the KNN regression finds *k* closest expression profiles in the available data and returns the mean of the appropriate targets. The D–GEX NNs were found to perform superiorly on all three datasets. The *L*_1_ and *L*_2_ regularized linear regression performed similarly to classical linear regression.

### 2.2 Adaptive activation functions

The activation function introduces non-linearities to neural networks and is crucial for networks’ performance. Usually, the same activation function is used for the whole network or at least for all neurons in a single layer even though this might be suboptimal. During the last decades, there were several attempts to use activation functions that might differ across neurons — e.g., [13]–[15]. The adaptive activation functions are getting more attention recently (e.g., [15]–[18]) and might become a new standard in the field. The simplest forms just add a parameter to a particular neural network that controls one of its properties (e.g., slope) while the more complex ones allow for learning of a large number of activation functions (e.g., adaptive spline activation functions in [14]).

However, adaptive activation functions (AAF) might be very useful even in the simplest form with a single added parameter — an AAF called *parametric rectified linear unit* (PReLU) was used for obtaining state-of-the-art result on the ImageNet Classification in 2015 first surpassing human-level performance [15]. The PReLU generalize the ReLU by adding parameter that controls the slope of the activation function for negative inputs (the ReLU is constant at zero for negative inputs) that is learned with other weights: *f* (*y*_*i*_) = *max*(0, *y*_*i*_) + *a*_*i*_*min*(0, *y*_*i*_), where *a*_*i*_ is the optimized parameter. The *leaky rectified linear unit* (LReLU) [19] is essentially a PReLU but with the parameter *a*_*i*_ fixed and not trainable. ELU (exponential linear unit) [20] is another popular activation function based on ReLU — it uses an exponential function for negative inputs instead of linear function. An adaptive ELU extension *parametric ELU* PELU together with mixing different activation functions using adaptive linear combination or hierarchical gated combination of activation function was shown to perform well [21].

A sigmoid activation function with shape autotuning [22] is an example of an early adaptive activation function. The generalized hyperbolic tangent [23] introduces two trainable parameters that control the scale of the activation function. A more general approach was introduced in [24] which used networks with a trainable amplitude of activation functions, the same approach was later used for recurrent neural networks [25]. A similar approach to trainable amplitude and generalized tanh is the so-called *neuron-adaptive activation function* (NAF) [26]. Other examples include networks with adaptive polynomial activation function [27], slope varying activation function [28] or back-propagation modification resulting in AAF [28], [29].

Another generalization of ReLU is the *adaptive piece-wise linear unit* (APLU) which uses the sum of hinge-shaped functions as the activation function [30]. An approach extending APLU is *smooth adaptive activation function* SAAF with piece-wise polynomial form and was specifically designed for regression and allows for bias–variance trade-off using a regularization term [17].

More complex approaches include spline interpolating activation functions (SAF) [14], [18], [31]–[33] which facilitate training of a wide variety of activation functions. A different approach represents the *network in network* (NIN) [34] which uses a micro neural network as an adaptive activation function. Another approach employs a gated linear combination of activation functions for each neuron which allows each neuron to choose which activation function (from an existing pool) it may use for minimizing the error [16]; a similar method uses just binary indicators instead of the gates [35]. The adaptive activation function might also be trained in a semi–supervised manner [36]–[38].

## 3 Methods

In order to examine whether our novel transformative adaptive activation function in the D–GEX model could lead to lower error, we have used the very same data as in [6].

### 3.1 Data

We have used gene expression data from the Affymetrix microarray platform curated by the Broad Institute that were provided by authors of the original D–GEX [6]. It contains 129,158 profiles each consisting of 22,268 probes. We have replicated the data pre–processing process presented in [6] — we have removed the biological and technical replicates and have used the same set of target and landmark genes. We have used 942 landmark genes to reconstruct the expression of 9,518 genes. The dataset was split into training, validation and testing set (the training set has 88,256 samples while the validation set has 18,895 samples and the testing set has 18,951 samples). The validation dataset was used for model selection and parameter optimization while the testing set was used for reporting the performance of selected models based on out-of-sample data.

### 3.2 Data normalization

The data were preprocessed in the same manner as in [6] except for the last step — the scaling to the interval to the zero mean and unit standard deviation. Scaling each variable separately as in [6], however, removes the absolute differences in expression between individual genes and it gives the same importance to all genes including the ones whose expression is near noise levels from the point of view of the error metrics. To keep the information about differences in expression levels, we scaled the data by transforming the whole data matrix to have zero mean and unit standard deviation without taking into account that there are different genes — thus the more expressed genes will be proportionately more expressed even after the scaling. We believe that such scaling is more suitable in this case as the minimization of the error metrics during the fitting phase gives relatively higher importance to more expressed genes and less to the genes whose expression is near the noise level. We have also verified that TAAF leads to performance gains in the original D–GEX also for the data scaled as in [6], details are available in the supplementary materials.

### 3.3 D–GEX

D–GEX, as proposed in [6], is a feedforward neural network consisting of one to three fully connected hidden layers each having the same number of neurons. The output layer consists of one neuron per target with a linear activation function. As in the original D–GEX, we have split the set 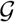 of 9,518 genes into two random subsets each containing half of the genes to enable learning on GPUs with smaller memory. A separate network was then trained using each of the sets and the final reconstruction consists of outputs of both networks. The original D–GEX used *dropout* [39] as a regularization technique to improve the generalization of the network with three different dropout rates — 0%, 10%, and 25%. Since the D–GEX with the 25% dropout rate had the best generalization [6], we have used only this rate for our experiments. All models were trained for 250 epochs (no improvement was observed near the end of the training), and the performance of the model from each epoch was evaluated on the validation data, only the best model from all epochs was used for further evaluation. The model optimization was done using the Nadam optimizer [40] with the learning rate *μ* = 0.0005 and optimizer specific parameters *β*_1_ = 0.9, *β*_2_ = 0.999, and schedule decay *η* = 0.004; the batch size was set to 256 profiles.

### 3.4 Model evaluation

To evaluate the model, we use the absolute error — first, we compute the *mean absolute error* of prediction MAE_*m*_(*s*) of model *m* for sample 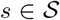 over individual genes 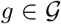 as in Equation (1) where *y*(*g, s*) is the expression of gene *g* for sample *s* and 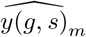 is the prediction of model *m* for the same target.

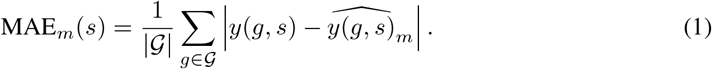

For further evaluation, we treat individual samples as independent (which is close enough to reality — our dataset probably contains small groups of samples which might be somewhat dependent, e.g. having same treatment, but it should be negligible for our size of dataset), thus for pairwise comparison we compare error metrics over individual samples and not over individual genes which have ties to each other. The overall performance of model *m* is defined as:

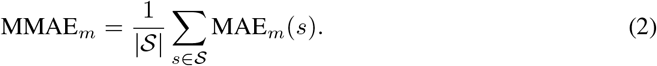

Pairs of models are not compared only in terms of MMAEs but also using pairwise differences. 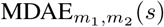 for models *m*_1_ and *m*_2_ and sample *s* is defined as:

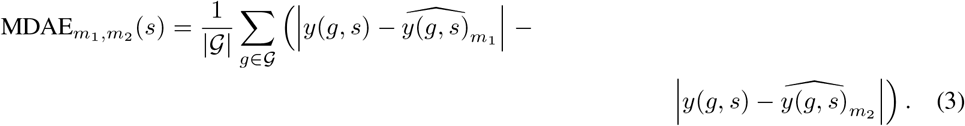

The *mean mean difference of absolute errors* 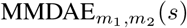 is defined as:

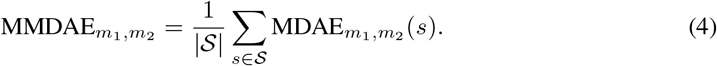

The MMDAE and its confidence interval that is estimated using bootstrap on MDAE allow to accurately compare two models even though their MMAEs are very close and their confidence intervals estimated using bootstrap on MAEs are overlapping.

To complement the model comparison based on MMDAEs, we have used Student’s paired t-test and paired Wilcoxon rank test on MAEs of individual samples. These tests were used to test the hypothesis that the differences in MAEs for individual samples over all genes are significantly different.

### 3.5 Proposed transformative adaptive activation function

We propose a novel family of adaptive activation functions to further improve the original D–GEX. The proposal is based on adaptive transformation of existing activation functions. The novel transformative adaptive activation function (TAAF) *g*(*f, y*) introduces four new parameters *α*, *β*, *γ*, 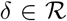 per neuron that transform the original activation function *f* (*y*) (called *inner activation function* in the context of the TAAF):

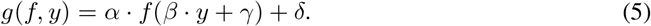

The output of a neuron with TAAF with inputs *x*_*i*_ is:

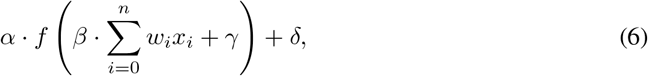

where *x*_*i*_ are individual inputs and *w*_*i*_ are its weights. If there is no unit *x*_*i*_ (i. e. unit constant), then the parameter *γ* is equivalent to the bias term of the neuron. The parameters are treated similarly as other weights in the NNs and are learned using back-propagation and gradient descent — the only differences is that parameters *α* and *β* are initialized to one and *γ* and *δ* are initialized to zero in every neuron.

The transforms allow for translation and scaling of the original activation function, and this transformation may be different for each neuron, i. e., each neuron has four additional parameters that define the TAAF for that neuron. The TAAF can also be viewed as a generalization of several existing adaptive activation functions — e.g. slope varying activation function [28] is the TAAF with adaptive parameter *β*, and frozen *α* = 1, and *γ*, *δ* = 0, or the trainable amplitude [24] is the TAAF with adaptive parameter *α*, and frozen *β* = 1, and *γ*, *δ* = 0. Other similar approaches also include parameters controlling slope but are focusing only on a special, predefined function [23], [26] instead of allowing any activation function to be used as the inner function in the TAAF.

#### 3.5.1 TAAF as output layer

It is a standard practice to use an output layer with a linear activation function as the sigmoidal activation functions such as hyperbolic tangent and logistic sigmoid have limited ranges. The original D–GEX architecture is no exception and uses a linear output layer. This, however, is no longer necessary with the use of TAAF as the scaling and translation allow for any interval range. The modified network architectures with TAAFs in the output layer (denoted TAAFo) enable better performance compared to the ones with a linear activation in the output layer by increasing the capacity of the network.

### 3.6 Implementation

The work was implemented in python 3, the neural networks were implemented using NN library *keras*^1^ [41] and computational framework *Tensorflow* [42]. Other used packages include *scipy* [43], *Scikit–learn* [44], *pandas* [45], and *NumPy* [46] for data manipulation and *matplotlib* [47] and *seaborn* [48] for visualizations.

## 4 Results

### 4.1 Experiment 1: D–GEX architectures

Preliminary experiments have shown that the *sigmoid* outperforms the original *tanh* function in the two–layered D–GEX architectures (details available in the Supplementary material). The next step was to establish the supremacy of the *sigmoid* function for other D–GEX architectures presented. The original D–GEX presented nine different architectures — one to three layers of the same size, each having either 3,000, 6,000, or 9,000 neurons. Since preliminary experiments have shown that the variance of the error for each repetition for both *sigmoid* and *tanh* activation function is very low, each architecture was trained only once due to computational resource constraints. As shown in Table 1, the *sigmoid* activation function dominates the *tanh* for all of the D–GEX architectures for both versions of the dataset. Table 1 shows the MMDAE between the sigmoid and tanh activation function — the 95% CI shows the improvement for all tested D–GEX architectures.

**Table 1:**
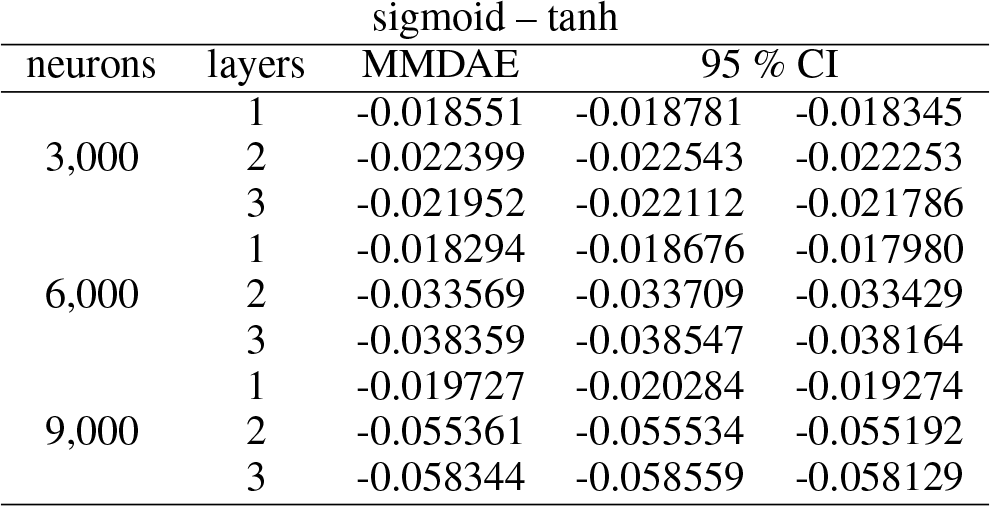
The MMDAE and its 95 % CI estimated using bootstrap on samplewise MDAEs for the sigmoid and tanh activation functions for D-GEX with 25% dropout on the test data.

| neurons | layers | sigmoid – tanh |  |  |
| --- | --- | --- | --- | --- |
|  |  | MMDAE | 95 % CI |  |
| 3,000 | 1 | -0.018551 | -0.018781 | -0.018345 |
|  | 2 | -0.022399 | -0.022543 | -0.022253 |
|  | 3 | -0.021952 | -0.022112 | -0.021786 |
| 6,000 | 1 | -0.018294 | -0.018676 | -0.017980 |
|  | 2 | -0.033569 | -0.033709 | -0.033429 |
|  | 3 | -0.038359 | -0.038547 | -0.038164 |
| 9,000 | 1 | -0.019727 | -0.020284 | -0.019274 |
|  | 2 | -0.055361 | -0.055534 | -0.055192 |
|  | 3 | -0.058344 | -0.058559 | -0.058129 |

### 4.2 Experiment 2: novel adaptive activation function

The goal of this and next experiments is to establish the improvement due to using the novel TAAFs. First, we compare the D–GEX architectures equipped with the sigmoid activation function to architectures equipped with the novel TAAF with sigmoid as the inner activation function. The results are shown in Table 2 where the models are compared using the MMDAEs. The table shows the signed difference in absolute errors between the classical sigmoid activation function and the adaptive activation function based on it — the novel transformative adaptive activation function was superior to classical sigmoid activation function for all D–GEX architectures. Furthermore, the means (medians) of MMAEs for both models were significantly different using Student’s paired t-test (Wilcoxon rank test) with *p* < 0.0001 for all tested D–GEX architectures.

**Table 2:** The MMDAE and its 95 % CI estimated using bootstrap on samplewise MDAEs for the TAAF with sigmoid as inner activation function and classical sigmoid activation function for D– GEX with 25% dropout on the test data.

| TAAF sigmoid – sigmoid |  |  |  |  |
| --- | --- | --- | --- | --- |
| neurons | layers | MMDAE | 95 % CI |  |
| 3,000 | 1 | -0.003434 | -0.003523 | -0.003330 |
|  | 2 | -0.003595 | -0.003636 | -0.003555 |
|  | 3 | -0.004464 | -0.004508 | -0.004419 |
| 6,000 | 1 | -0.004600 | -0.004748 | -0.004401 |
|  | 2 | -0.003362 | -0.003407 | -0.003315 |
|  | 3 | -0.005617 | -0.005674 | -0.005563 |
| 9,000 | 1 | -0.005145 | -0.005360 | -0.004860 |
|  | 2 | -0.004546 | -0.004598 | -0.004491 |
|  | 3 | -0.006799 | -0.006876 | -0.006724 |

However, such comparison does not suffice as the four added parameters per neuron increase the capacity of the neural network and the improvement might just be caused by the increase in the capacity. Indeed, it seems that increased capacity helps the D–GEX as the architectures with more neurons have lower prediction error for the same number of layers. We have reduced the number of neurons in each layer in the D–GEX with the TAAFs such that the total number of parameters is the same as in the original D–GEX with the same architecture. The number of removed neurons was always lower than 30 as the number of added weight per neuron is insignificant compared to the number of weights per incoming connections. The improvement of the reduced D-GEX with TAAFs was from 0.0034 to 0.0068 across different D-GEX architectures. The whole comparison with the reduced D–GEXs with the adaptive activation function based on sigmoid and the original D-GEXs is shown in the Supplementary materials. The models with the adaptive activation function are still superior to the original D–GEX — the reduction in the number of neurons was too small to result in a significant decrease in performance.

### 4.3 Experiment 3: importance of individual parameters

The proposed TAAF introduces four additional parameters, and so far we have not established the importance of individual parameters. Since the TAAF can be viewed as a generalization of several already established AAFs, the performance increase compared to the sigmoid activation function might be due only to those parameters that were already established as beneficial — e.g., trainable amplitude [24]. Furthermore, since the proposed adaptive activation function is applied to the weighted sum of inputs in the neuron, the parameter *β* might seem to be redundant:

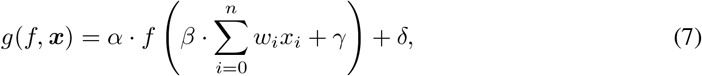

where *n* is number of inputs in the neuron, *x*_*i*_ are individual inputs and *w*_*i*_ are associated weights. This can be expressed without the parameter *β* if we define *u*_*i*_ = *βw*_*i*_:

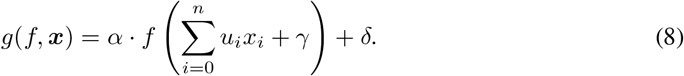

To verify that all TAAF parameters are improving the performance, we have trained neural networks with reduced TAAFs that had some of the parameters removed. We have evaluated all 16 subsets of TAAF parameters (from the reduced TAAF equivalent to classical sigmoid activation function to full TAAF with all four adaptive parameters) using three layerd D–GEX architecture with 6000 neurons in each layer. The networks with different subsets of TAAF parameters were pairwise evaluated based on MMDAE. The Figure 1a shows the MMDAEs between all model pairs while Figure 1b shows whether model A (row) is significantly better than model B (column) based on paired Wilcoxon rank test on samplewise MAEs at significance level *α* = 0.001. The full TAAF is significantly better than all other combinations of parameters which shows that the proposed TAAF with four parameters is the correct choice and that it outperforms other adaptive activation functions it generalizes.

**Figure 1:**
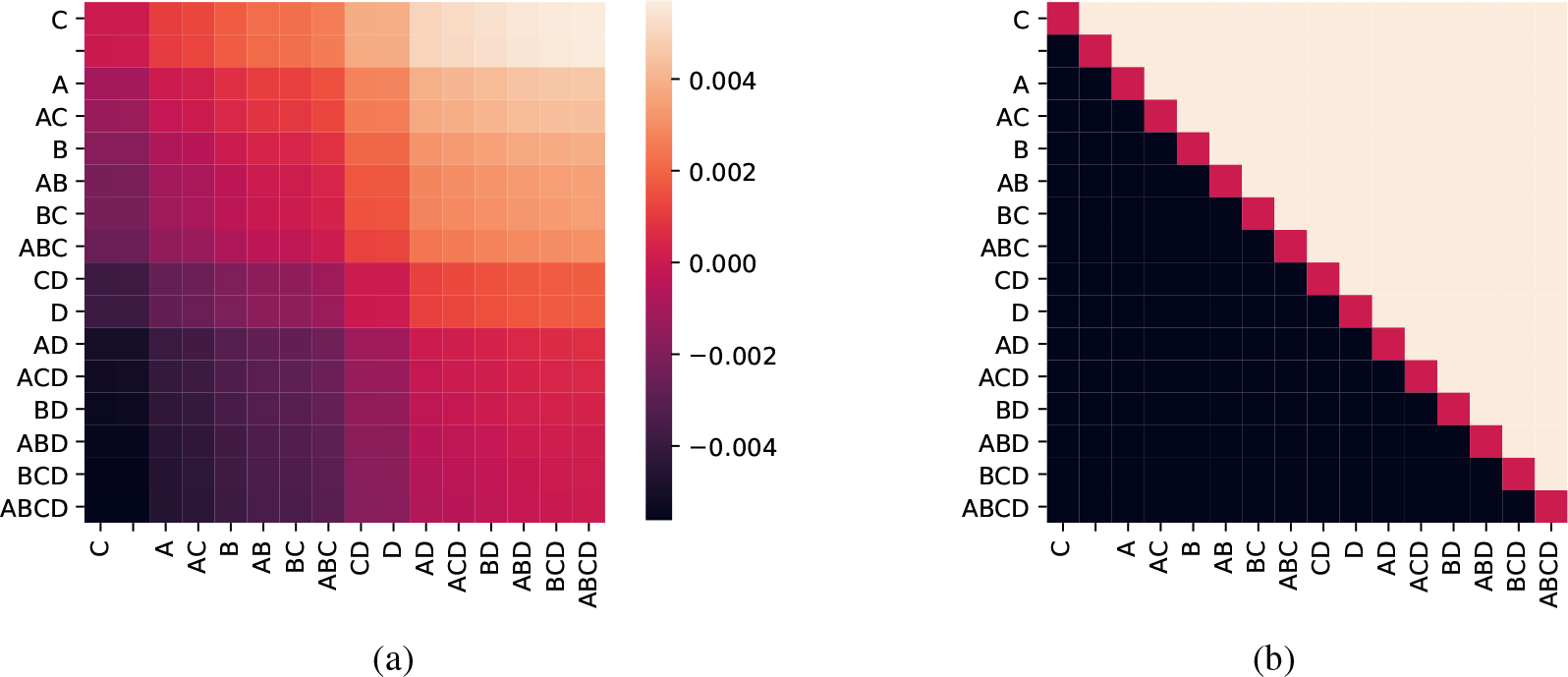
Evaluation of TAAFs with all possible subsets of adaptive parameters on out-of-sample data. Figure 1a shows the MMDAE while Figure 1b shows whether the prediction has lower MAE based on the paired Wilcoxon rank test at significance level 0.001. The model labels specify which adaptive parameters were used in the TAAF — e.g. `AD` means adaptive parameters *α* and *δ* were used.

### 4.4 Experiment 4 — TAAF in the output layer

The networks with TAAF do not require that the output layer contains a linear activation function for regression tasks as the TAAF allows for scaling and translation. Using TAAFs in the output layer might lead to better performance as shown in Table 3 where networks with TAAFs in the output layer are compared with networks with a linear output layer. The usage of TAAFs in the output layer was beneficial for all tested architectures.

**Table 3:** The MMDAE and its 95 % CI estimated using bootstrap on samplewise MDAEs for the TAAF with sigmoid as inner activation function and 25% dropout on the test data. `TAAFo sigmoid` denotes a network that contains TAAFs in the output layer while `TAAF sigmoid` uses a linear activation in the output layer.

| TAAFo sigmoid – TAAF sigmoid |  |  |  |  |
| --- | --- | --- | --- | --- |
| neurons | layers | MMDAE | 95 % CI |  |
| 3,000 | 1 | -0.000401 | -0.000472 | -0.000308 |
|  | 2 | -0.001015 | -0.001091 | -0.000945 |
|  | 3 | -0.001896 | -0.001951 | -0.001843 |
| 6,000 | 1 | -0.000679 | -0.000789 | -0.000531 |
|  | 2 | -0.001654 | -0.001718 | -0.001591 |
|  | 3 | -0.002474 | -0.002521 | -0.002428 |
| 9,000 | 1 | -0.000919 | -0.001075 | -0.000711 |
|  | 2 | -0.001864 | -0.001935 | -0.001796 |
|  | 3 | -0.001426 | -0.001477 | -0.001377 |

## 5 Overall comparison

The best single network performs much better than our reimplementation of the original D–GEX — the MAE of the best network (3*x*9, 000 TAAFo with sigmoid) is 0.1340 (the 95% CI estimated over samples is [0.13316, 0.13486]) compared to D–GEX with tanh activation function with MAE of 0.1637 (95% CI [0.16279, 0.16458]). Our proposed network performs better for 18,849 (99.75%) samples while worse only for 2 (0.001%) samples when the MAEs over genes for individual samples are compared using pairwise Wilcoxon rank test with significance level *α* = 0.0001.

All improvements to the original D–GEX are depicted in Figure 2a which shows the improvement of individual modifications (details about ensembling available in Supplementary materials). Figure 2b summarizes the individual improvements over the basic D–GEX with our proposed activation function. The Table 4 shows the absolute performance of the top ten D–GEXs.

**Table 4:** The 10 best D–GEX architectures in terms of MMAE on the test datat. 25% dropout was used.

| rank | MMAE | neurons | layers | type | activation |
| --- | --- | --- | --- | --- | --- |
| 1 | 0.134015 | 9,000 | 3 | TAAFo | sigmoid |
| 2 | 0.134503 | 9,000 | 2 | TAAFo | sigmoid |
| 3 | 0.135430 | 8,997 | 3 | TAAF (reduced) | sigmoid |
| 4 | 0.135442 | 9,000 | 3 | TAAF | sigmoid |
| 5 | 0.136064 | 9,000 | 2 | TAAFo | tanh |
| 6 | 0.136367 | 9,000 | 2 | TAAF | sigmoid |
| 7 | 0.136774 | 8,997 | 2 | TAAF (reduced) | sigmoid |
| 8 | 0.136883 | 9,000 | 3 | TAAFo | tanh |
| 9 | 0.137154 | 9,000 | 3 | TAAF | sigmoid |
| 10 | 0.137189 | 6,000 | 3 | TAAFo | sigmoid |

**Figure 2:**
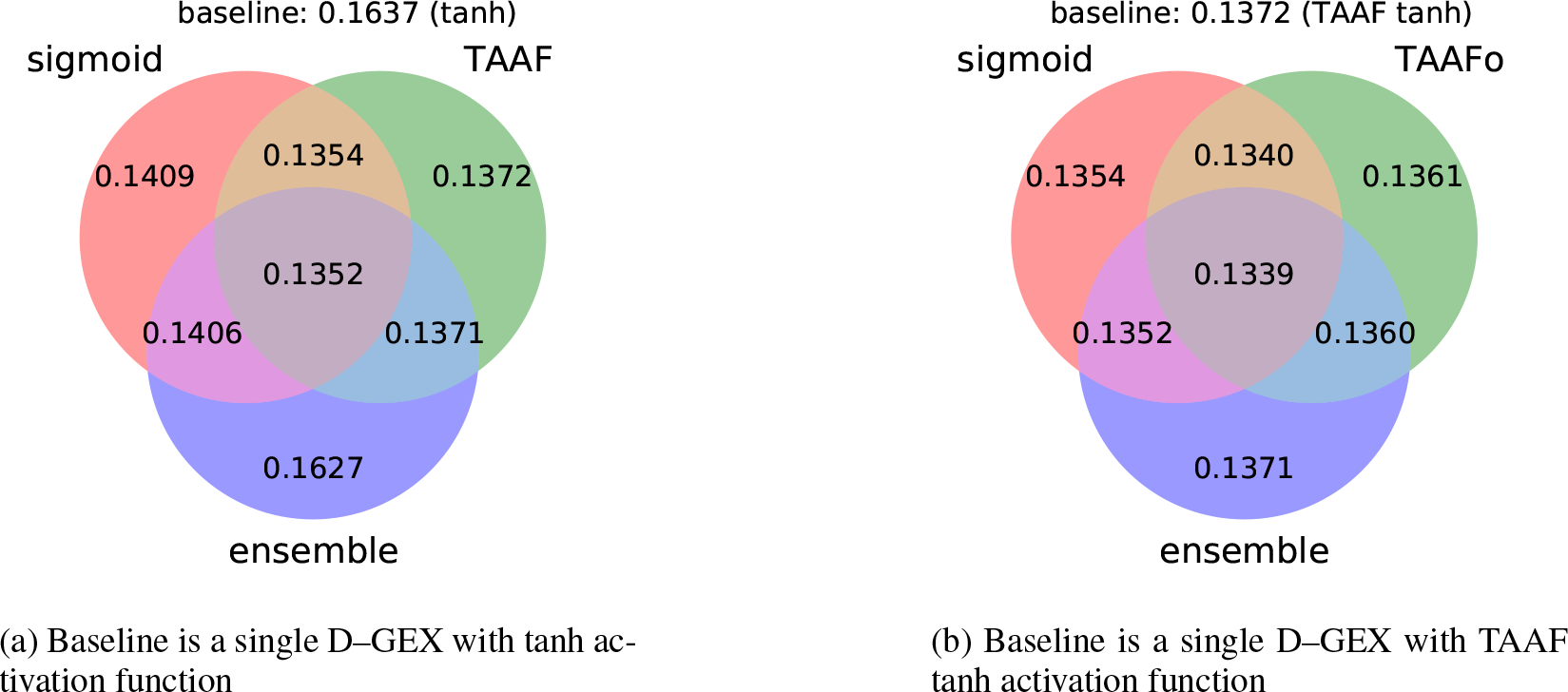
The Venn diagram depicting the performance for individual improvements over the standard D–GEX baseline with tanh activation function. The diagram shows the best MMAE over all D–GEX architectures for given approach.

## 6 Conclusion

We have evaluated the use of different activation functions in the D–GEX architectures and found that the hyperbolic tangent function that was used on the original D–GEX is not the most suitable one. We have evaluated five other activation functions on one half of the target genes (4,759 genes) in a ten times repeated experiment for a single D–GEX architecture and have found that the sigmoid activation function outperforms the others. This result was confirmed for all nine presented D–GEX architectures on two different versions of the dataset. We conclude that the sigmoid activation functions are the better choice for this task despite the tanh being usually more suitable and recommended in [49].

Furthermore, we proposed a novel family of transformative adaptive activation functions (TAAFs) which allows for scaling and translation of any other activation function. We have evaluated the novel TAAF in the D–GEX settings and verified that it outperforms its non-adaptive counterparts. The proposed TAAF generalizes several existing adaptive activation functions [24], [28] and performs superiorly. Our single selected network significantly outperforms the best D–GEX architecture with *p* < 0.0001 and improves the MAE by ≈ 18%.

## Acknowledgements

We gratefully acknowledge the support of NVIDIA Corporation with the donation of the Titan Xp GPU used for this research. Additional computational resources were supplied by the Ministry of Education, Youth and Sports of the Czech Republic under the Projects CESNET (Project No. LM2015042) and CERIT-Scientific Cloud (Project No. LM2015085) provided within the program Projects of Large Research, Development and Innovations Infrastructures.

## Funding

This work was supported by the Grant Agency of the Czech Technical University in Prague, grant No. SGS17/189/OHK3/3T/13. The work was supported by the grant 17-31398A from the Ministry of Health of the Czech Republic.

## Footnotes

1 https://keras.io

